# An updated inventory of genes essential for oxidative phosphorylation identifies a mitochondrial origin in familial Ménière’s disease

**DOI:** 10.1101/2025.01.29.635272

**Authors:** Marcell Harhai, Mads M. Foged, Emeline Recazens, Miriam Lisci, Sylvain Chollet, Juan C. Landoni, Nora Laban, Suliana Manley, Jose Antonio Lopez-Escamez, Anna Lysakowski, Alexis A. Jourdain

## Abstract

Mitochondrial disorders (MDs) are among the most common inborn errors of metabolism and primarily arise from defects in oxidative phosphorylation (OXPHOS). Their complex mode of inheritance and diverse clinical presentations render the diagnosis of MDs challenging and, to date, most lack a cure. Here, we build on previous efforts to discover genes necessary for OXPHOS and report a highly complementary galactose-sensitized CRISPR-Cas9 “growth” screen, presenting an updated inventory now with 481 OXPHOS genes, including 157 linked to MDs. We further focus on *FAM136A*, a gene associated with Ménière’s disease, and show that it supports inter-membrane space protein homeostasis and OXPHOS in cell lines, mice, and patients. Our study identifies a mitochondrial basis in a familial form of Ménière’s disease (fMD), provides a comprehensive resource of OXPHOS-related genes, and sheds light on the pathways involved in mitochondrial disorders, with the potential to guide future diagnostics and treatments for MDs.

**Graphical abstract:** 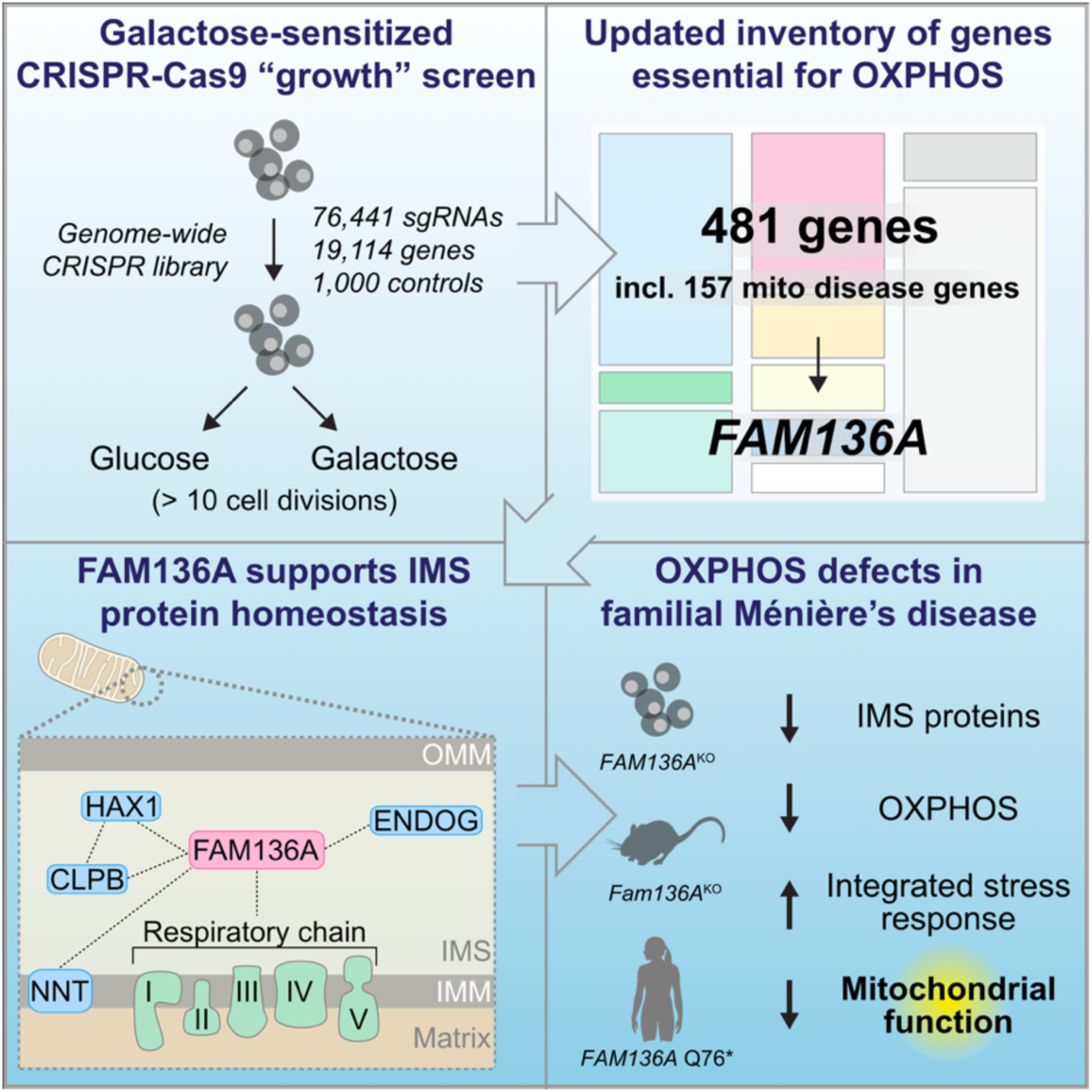

**Bullet points:**

- Genome-wide CRISPR-Cas9 growth screening complements death screening
- 481 genes essential for OXPHOS, including 157 mitochondrial disease genes
- *FAM136A* supports mitochondrial intermembrane space protein homeostasis
- Depletion of *FAM136A* in Ménière’s disease models causes OXPHOS defects

## Introduction

Mitochondrial disorders (MDs) are a large, heterogeneous group of diseases. They are the most prevalent inborn errors of metabolism, affecting 1 in 5000 people, and their manifestations may target organs in either a multisystemic or a tissue-specific manner^1,2^. The onset of MDs spans all ages, from newborns to the elderly, with diverse clinical and tissue manifestations, including lactic acidosis, myopathies, encephalopathy, neurodegeneration and peripheral neuropathies, diabetes mellitus, and deafness^2,3^. MD primarily arises from defects in mitochondrial oxidative phosphorylation (OXPHOS), which requires both the nuclear and mitochondrial genomes. To date, more than 300 genes from both genomes have been linked to MDs in patients^4^. With such a complex genetic background, pathophysiology, and clinical manifestation, MDs pose a great challenge in diagnosis. The currently available diagnostic methods combining clinical assessment, tissue analysis, and genetic screens yield suboptimal diagnostic rates, and molecular genetics analysis fails to identify the causative genetic mutation in ∼50% of cases^5–7^. To date, most MDs lack a cure, and further research is needed to increase our understanding of mitochondrial functions, identify novel MD genes, and develop appropriate treatments.

To identify genes essential for OXPHOS, whose mutations could cause MDs, we and colleagues previously performed a nutrient-sensitized genome-wide CRISPR-Cas9 screen based on glucose auxotrophy^8^, a phenotype by which MD suspicion may be confirmed through lack of proliferation in glucose-free, galactose-containing medium^9,10^. We found that cells with mitochondrial dysfunction actively die in galactose medium and developed a protocol of “death screening” whereby dead cells are selected and single-guide RNAs (sgRNAs) are sequenced. Death screening effectively converts negative selection “drop out” screens into positive selection screens, and we recently reported a refined death screening protocol called Dead-Seq^11^. With this first galactose-sensitized screen, we built an inventory of 300 genes essential for OXPHOS in K562 myelogenous leukemia cells, representing several mitochondrial and non-mitochondrial pathways and including 71 “positive control” genes previously associated with MDs. Our work highlighted how CRISPR-Cas9 screening is a powerful tool to improve the discovery of genes involved in MDs. However, while death screening was performed with high precision, it seemed to lack sensitivity as many known MD genes were not recovered. A plausible explanation is the acute nature of death screening, which was performed after only 24 hours of incubation in a glucose- or galactose-containing medium (**Figure 1A**). Reminiscent of late-onset MDs, genes and pathways with milder or longer-term impact on mitochondria may have been missed.

**Figure 1:**
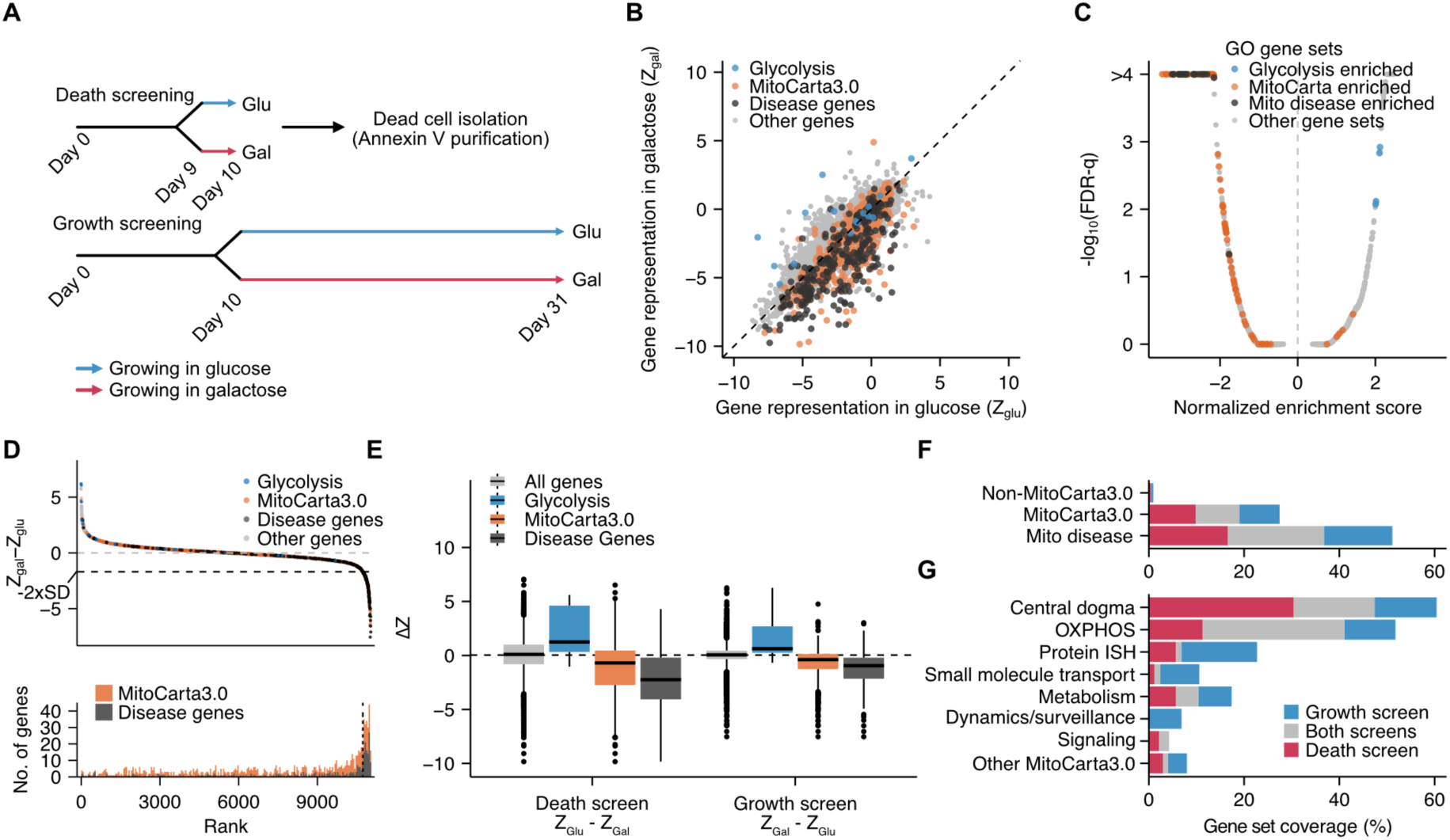
Genome-wide CRISPR-Cas9 galactose-sensitized screens identify genes essential for OXPHOS. **(A)** Schematic overview of death and growth screening. Glu: 25 mM glucose. Gal: 25 mM galactose. **(B)** Gene representation of the genome-wide CRISPR-Cas9 growth screen results in glucose vs galactose showing Z scores relative to non-targeting controls in glucose (Z_Glu_) and galactose (Z_Gal_). N = 11,020 expressed genes. **(C)** GSEA highlighting pathways identified in our genome-wide CRISPR-Cas9 growth screens in glucose vs galactose screen (ΔZ = Z_gal_ – Z_glu_). Each dot represents a gene ontology gene set. Orange: gene sets with more than 50% MitoCarta3.0 genes. Blue: gene sets with more than 50 % of the genes being canonical glycolysis enzymes. Dark grey: gene sets with more than 50 % mitochondrial disease genes. **(D)** Ranked growth screen ΔZ scores. Each point represents an individual gene. Genes involved in glycolysis (blue), as well as MitoCarta3.0 genes (orange) and MD genes (dark grey), are highlighted. Black dashed line indicates the threshold (−2xSD of ΔZ scores) for defining “hits” in the screen. Lower panel shows the ranked abundance of MitoCarta3.0 genes (orange) and MD genes (black) as they appear in the upper panel. **(E)** Pathway comparison between growth and death screening. Boxes indicate the interquartile range (IQR) Points represent outliers (outside 1.5 times the IQR above the 0.75 quantile or below the 0.25 quantile). **(F-G)** Cumulative analysis of gene recovery in death and growth screening expressed as a percentage of the indicated gene sets. Gene sets in G are the 7 top-level MitoPathways. ISH: import, sorting, and homeostasis.

Here, we present a nutrient-sensitized CRISPR-Cas9 screen based on the proliferation of cells in a galactose medium over 21 days. This “growth” screening protocol exhibits strong complementarity to death screening, and we report an updated inventory of genes essential for OXPHOS, now including 157 positive control MD genes. Among the new genes and pathways discovered, we uncover that *FAM136A*, a poorly characterized gene associated with Ménière’s disease, a chronic inner ear disorder defined by episodes of vertigo, sensorineural hearing loss and tinnitus^12^, supports mitochondrial intermembrane space (IMS) protein homeostasis and OXPHOS across several models. Our data reveal a likely mitochondrial dysfunction involved in familial Ménière’s disease pathophysiology, and more generally highlights the power of genetic screening in discovering human disease genes.

## Results

### Galactose-sensitized CRISPR-Cas9 growth screening complements death screening

To complement our death screen, we designed a “growth screen” based on the proliferation of cells in glucose or galactose over several weeks (**Figure 1A**). We employed the Brunello lentiviral genome-wide CRISPR-Cas9 library with 76,441 single-guide RNAs (sgRNAs) to systematically target 19,114 distinct human genes and 1,000 controls in K562 cells^13,14^. We infected cells in a glucose-rich medium supplemented with uridine and pyruvate required for cell growth with OXPHOS defects^15,16^. Following 10 days of growth to ensure CRISPR-Cas9-mediated gene knockout, we swapped to either glucose- or galactose-containing medium. After another 21 days, corresponding to > 10 cell divisions, we harvested the cells and performed next-generation sequencing to determine the abundance of sgRNAs in each condition (**Table S1A**). We analyzed the screen using a method based on Z-scores^17^ and gene set enrichment analysis (GSEA)^16,18^. As expected, we found that sgRNAs targeting genes encoding mitochondrial proteins (MitoCarta3.0^19^) and MD genes were significantly depleted in galactose (**Figure 1B-C**), whereas genes involved in glycolysis were generally dispensable in galactose^20,21^. At a threshold of two standard deviations (SD) (**Figure 1D**), our growth screen identified 291 genes, corresponding to 200 MitoCarta3.0 proteins, including 106 MD genes, and 91 non-MitoCarta3.0 genes (**Figure S1A, Table S1B**).

Comparing our growth screen to our death screen^8^, we found that both identified non-MitoCarta3.0, MitoCarta3.0, and MD genes at similar rates, with only a partial overlap highlighting complementarity (**Figure 1E-F, S1B**). To better understand these differences, we classified hits from both screens into the seven major MitoCarta3.0 pathways. We observed that (I) both screening modalities identified “OXPHOS” and “metabolism” genes equally well, with “OXPHOS” being the category showing the highest overlap, (II) death screening performed better at identifying genes involved in mitochondrial gene expression (“central dogma” genes), and (III) growth screening outperformed death screening at identifying genes involved in “mitochondrial protein import, sorting, and homeostasis”, “small molecule transport” and “mitochondrial dynamics and surveillance” (**Figure 1G, Table S2A**). Looking at genes encoding non-mitochondrial proteins, we found that some were part of pathways previously highlighted with death screening, such as AMPK signaling and the INO80 chromatin remodeling complex, while others were identified only with growth screening. To further validate these new genes, we used CRISPR/Cas9 to deplete the expression of 8 hits from the screen: the cytosolic arm of heme metabolism (*ALAD*, *HMBS*, *UROS*), the plasma membrane metal transporters *SLC31A1* (copper transport) and *SLC40A1* (iron transport), the cholesterol biosynthesis gene *SQLE*, the Poly(A) polymerase alpha subunit *PAPOLA*, as well as the cytosolic RNA-binding protein *CLUH* involved in mitochondrial biogenesis^22^ as control. To assess respiratory chain integrity, we quantified 6 proteins of the mitochondrial respiratory complexes by immunoblotting and observed a significant decrease in at least one respiratory chain subunit in 6 of the 8 genes tested (**Figure S1C-D**). Our data highlights how these screening strategies complement each other and synergistically support the identification of genes essential for OXPHOS.

### An updated inventory of human genes required for oxidative phosphorylation

Our observations on the complementarity between the two screening modalities prompted us to combine their data and generate an updated inventory of the genes required for oxidative phosphorylation in humans (**Figure 2, Table S2A**). Our updated inventory now comprises 481 genes, including 314 MitoCarta3.0 genes, among them 157 MD genes, as well as 167 non-MitoCarta3.0 genes. We organized genes based on the seven major MitoCarta3.0 pathways, with “central dogma” and “OXPHOS” being the most represented pathways (**Figures 1G, 2, and S2**) and calculated a confidence score for each gene based on ranks in the screens. We further compiled a collection of 1,145 sgRNA sequences corresponding to the two best sgRNAs for each gene hit in our screens, useful for future mitochondrial research (**Table S2B**). Our updated inventory also includes eight genes not currently assigned to a MitoCarta3.0 pathway, among which we noticed *FAM136A*, previously reported to encode an uncharacterized mitochondrial protein whose mutations are associated with a familial form of Ménière’s disease^23^ (fMD).

**Figure 2:**
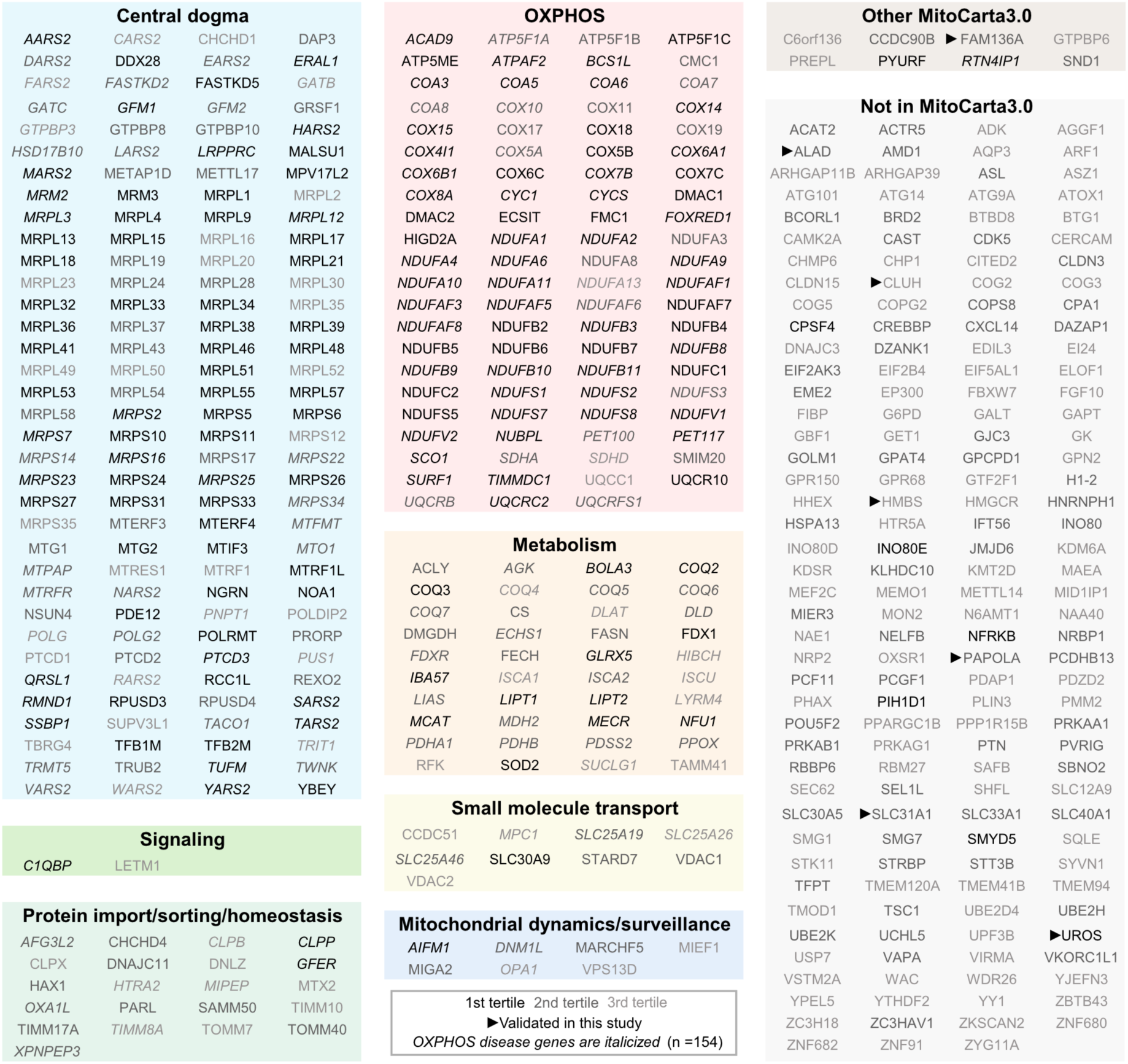
Updated inventory of human genes essential for OXPHOS. The inventory combines death and growth screening and now comprises 481 genes organized by mitochondrial pathway, with 314 MitoCarta3.0 and 167 non-MitoCarta3.0 genes (see Methods). MitoCarta3.0 genes are further divided into the 7 top-level MitoPathways^19^. Italics: 157 genes involved in MDs. Genes are shaded according to their confidence scores (see Methods) from high confidence (1st tertile) to lower confidence (3rd tertile). Arrowheads: genes investigated in this study. HAX1 and GTPBP8 are not represented in MitoCarta3.0 and were manually assigned based on the previous studies^45,46^. FAM136A is found under “Other MitoCarta3.0”.

### Ménière’s disease-associated gene *FAM136A* is required for inter-membrane space protein homeostasis

Ménière’s disease (idiopathic endolymphatic hydrops) is an inner ear condition characterized by vertigo, hearing loss, and tinnitus. The presence of *FAM136A* in our inventory was unexpected because Ménière’s disease is not considered to be an MD, even though it shares symptoms with some MDs. FAM136A is a eukaryote-specific mitochondrial inter-membrane space protein of unknown function^19,24^. It is predicted to be composed of three α-helices, and the disease-associated mutation (GRCh38 chr2:70300842G>A) introduces a premature stop codon and truncates the protein from the middle of the second helix (FAM136A-Q76*), likely disrupting its function and localization (**Figure 3A**). To investigate the potential mislocalization of FAM136A-Q76*, we expressed FLAG-tagged cDNAs corresponding to the wild-type and patient (FAM136A_1-75_) protein in U-2 OS cells and performed super-resolution 3D instant structured illumination microscopy (iSIM) imaging. While wild-type FAM136A-FLAG localized to mitochondria, we only achieved low expression of FAM136A_1-75_-FLAG, which was mislocalized to the cytosol (**Figure 3B**) and showed a dramatic reduction in half-life (**Figure 3C**). These results strongly suggest that disease-associated FAM136A-Q76* is a loss-of-function mutation, encoding an unstable and mislocalized protein.

**Figure 3:**
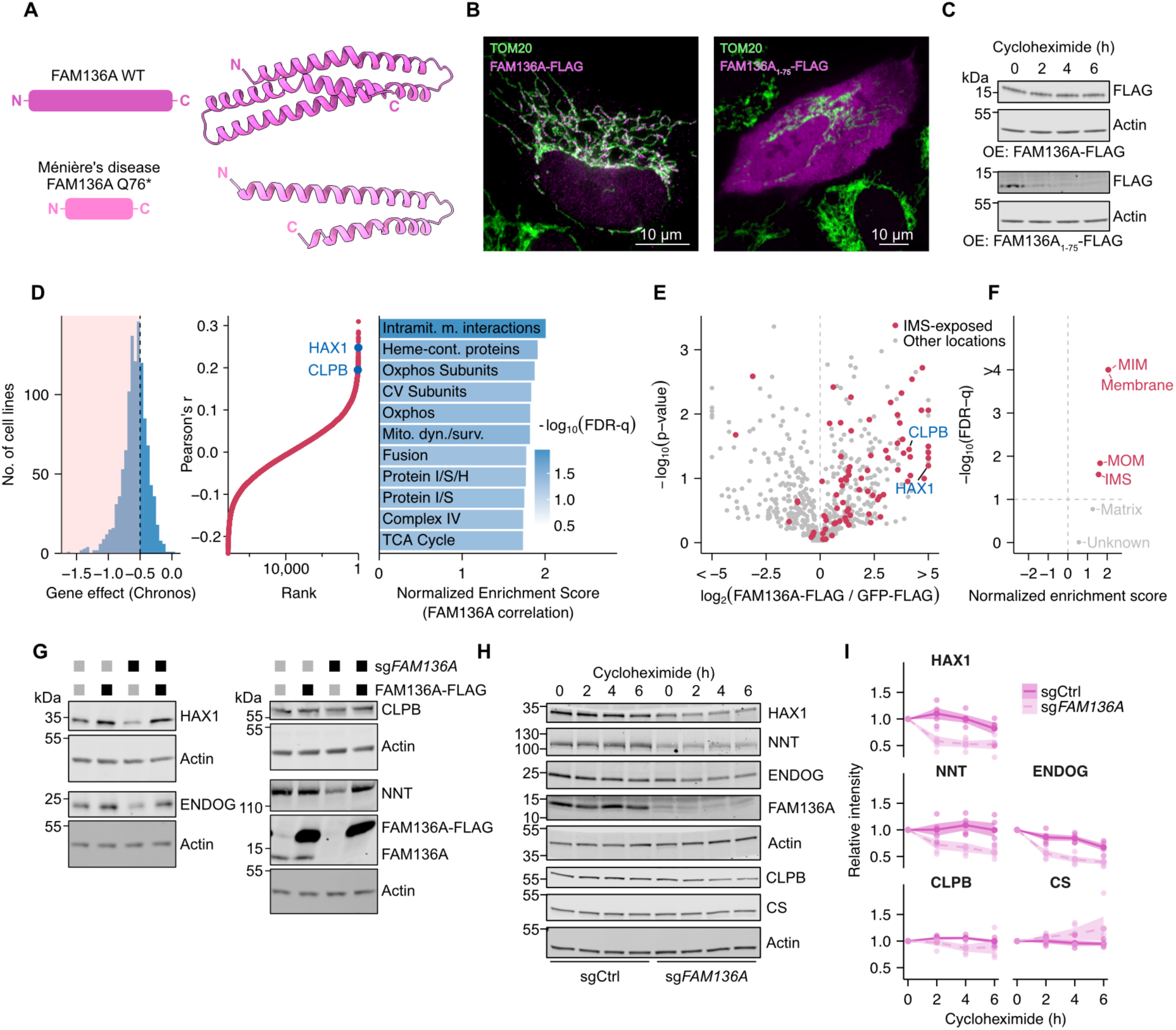
FAM136A supports IMS proteostasis. **(A)** Predicted structures of wild-type (WT) FAM136A and Meniere’s disease-associated mutant FAM136A-Q76*. **(B)** Immunofluorescence of K562 cells over-expressing FAM136A-3xFLAG or FAM136A_1-75_ in U2-OS cells with antibodies targeting FLAG and the outer membrane protein TOM20. **(C)** Temporal protein stability assay of overexpressed WT FAM136A compared to overexpressed (OE) FAM136A_1-75_. **(D)** Analysis of *FAM136A* CCLE gene effect. Left panel: distribution of gene effect Chronos scores for each of the 1,150 CCLE cell lines, highlighting the scores below −0.5. Middle panel: gene-gene Pearson correlation coefficients between FAM136A and other genes in the CCLE ranked by correlation coefficients. HAX1 and CLPB are highlighted in blue. Right panel: MitoPathways GSEA of the FAM136A correlation coefficients showing pathways with FDR-q < 0.05. Intramit. m. interactions: Intramitochondrial membrane interactions; Heme-cont. proteins: Heme-containing proteins; Mito. Dyn./surv.: Mitochondrial dynamics and surveillance; Protein I/S/H: protein import, sorting, and homeostasis; Protein I/S: protein import and sorting. FDR: False discovery rate. **(E)** FLAG IP-MS of mitochondria isolated from K562 cells overexpressing either FAM136A-FLAG or GFP-FLAG. Intermembrane-space (IMS)-exposed proteins, defined as proteins annotated as either “inner membrane”, “outer membrane”, “membrane” or “IMS” in the MitoCarta3.0 database^19^, are highlighted in red. **(F)** Gene set enrichment analysis of the IP-MS data shown in E. IMS-exposed pathways are highlighted in red. **(G)** Immunoblotting of IMS-localized proteins in control (sgCtrl) vs *FAM136A*-depleted (sg*FAM136A*) K562 cells expressing GFP-FLAG or FAM136A-FLAG. **(H)** Temporal protein stability assay of IMS-exposed proteins in sgCtrl and sg*FAM136A* K562 cell lines. **(I)** Quantification of H. Shaded areas represent mean values +/- SEM. n = 4 replicates.

To further understand the function of *FAM136A,* we analyzed its expression in publicly available databases^25^. Similar to OXPHOS genes, we found *FAM136A* to be expressed in all tissues as well as in most cancer cells and is not limited to the inner ear (**Figure S3A-B**). We further analyzed the genetic dependency of *FAM136A* across 1,150 cancer cell lines from the Cancer Cell Line Encyclopedia (CCLE) and found that cells are selectively dependent on *FAM136A* (CCLE Chronos^26^ score < −0.5 in 66.3% of the cell lines), irrespective of their lineages (**Figure 3D, S3C**). Comparing the *FAM136A*-dependency profile across the 1,150 cancer cell lines with the dependency profiles of 17,930 genes and using MitoPathways, we found that *FAM136A* correlated best with genes involved in pathways such as “intramitochondrial membrane organization”, “protein import, sorting, and homeostasis” and “OXPHOS” (**Figure 3D**) together hinting at a role for *FAM136A* in IMS protein homeostasis (proteostasis).

Next, we stably expressed FAM136A-FLAG in K562 cells and performed immunoprecipitation and mass spectrometry (MS) to discover its interacting proteins. We observed a significant enrichment of IMS-exposed proteins (**Figure 3E-F, Table S3**), an observation that is corroborated by a published crosslinking and MS analysis of whole mitochondria^24^. Among these IMS-exposed proteins, we noticed a co-essentiality pattern with CLPB, an IMS protein disaggregase, and HAX1, an intrinsically disorganized protein and substrate of CLPB^27,28^ (**Figure 3D**), suggesting a functional connection. Accordingly, we observed a marked reduction of both proteins in *FAM136A*-depleted cells (**Figure 3G**), which was rescued by introducing FAM136A-FLAG cDNA. We further tested whether additional IMS proteins were depleted in the absence of *FAM136A*. Testing for a soluble and a membrane protein, we observed decreased levels of the endonuclease G ENDOG and the NAD(P) transhydrogenase NNT, respectively, indicating that the effect of *FAM136A* depletion expands to multiple substrates in the IMS (**Figure 3G**). To test whether the decrease in abundance was related to protein stability, we next blocked cytosolic protein synthesis using cycloheximide and observed a lower half-life of IMS HAX1, ENDOG, NNT, and to a lesser extent CLPB, but not matrix citrate synthase (CS), indicating a destabilizing effect of *FAM136A* depletion on these IMS proteins (**Figure 3H-I**). Taken together, we conclude that FAM136A is required for mitochondrial IMS proteostasis.

### *FAM136A* supports OXPHOS in cell lines, mice, and patients

We next investigated the possibility of a global role of *FAM136A* in IMS proteostasis and performed total cell proteomics on *FAM136A*-depleted K562 cells (**Table S4A**). We confirmed the depletion of HAX1, CLPB, ENDOG, and NNT **(Figure 3G-I, S4A)**, as well as proteins from the respiratory chain, which is partly exposed to the IMS (**Figure 4A-B).** We also noticed a decrease in mitochondrial central dogma proteins, but no effect on other mitochondrial pathways, indicating no global depletion of mitochondrial content (**Figure 4A-B)**. We validated decreased levels of respiratory chain proteins, including NDUFB8 and MT-CO1 (**Figure 4C**), alongside a reduction in cellular respiration (**Figure 4D**). The effect of *FAM136A* depletion on the respiratory chain complexes and respiration validates our CRISPR-Cas9 screen.

**Figure 4:**
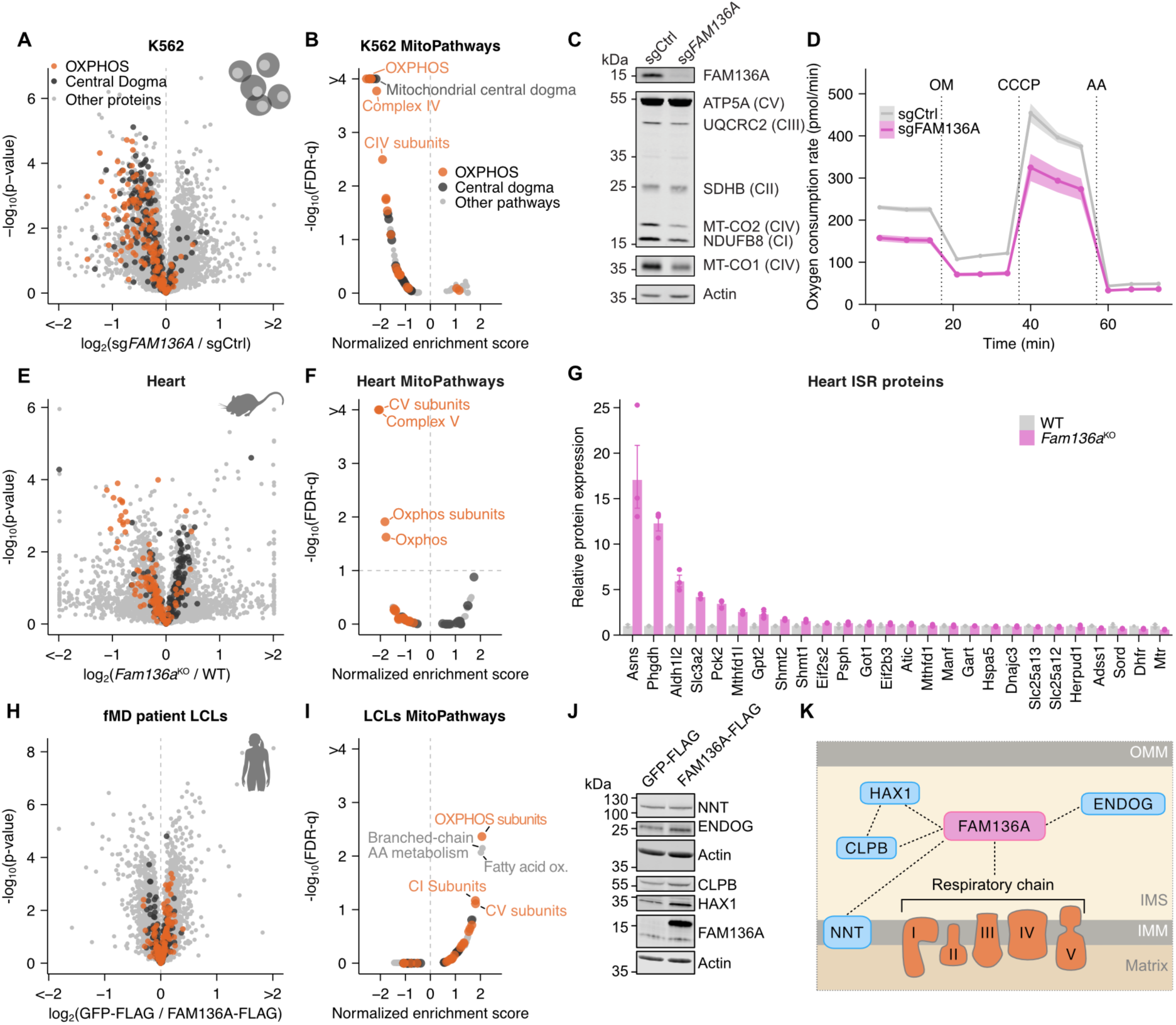
FAM136A supports mitochondrial oxidative phosphorylation. **(A)** Quantitative proteomics comparing FAM136A-depleted (sg*FAM136A*) to control (sgCtrl) K562 cell lines. n = 4 independent viral infections per condition. Proteins involved in mitochondrial central dogma (black) and OXPHOS (orange), as defined in MitoCarta3.0, are highlighted. **(B)** MitoPathways GSEA of K562 proteomics data (Figure 4A). **(C)** Immunoblot of sg*FAM136* and control K562 cell lines showing FAM136A and OXPHOS subunits. **(D)** Oxygen consumption rates in sg*FAM136* vs control K562 cell lines. OM: Oligomycin; AA: Antimycin A, CCCP: carbonyl cyanide m-chlorophenylhydrazone. n = 4 independent viral infections. Shaded areas indicate mean values +/- SEM. **(E)** Quantitative proteomics comparing protein levels in heart tissue of *Fam136a*^KO^ vs wild type (WT) mice. n = 3 mice per condition. **(F)** MitoPathways GSEA of mouse heart proteomics data in E. **(G)** Bar plots showing log_2_(fold changes) of proteins involved in the ISR in the mouse heart proteomics datasets. n = 3 mice per condition. ISR genes were manually assigned based on previous studies^30–33^ (**Table S4F**). **(H)** Quantitative proteomics comparing protein levels in fMD patient LCLs with a heterozygous FAM136A-Q76* mutation overexpressing either FAM136A-FLAG or GFP-FLAG. n = 4 replicates per condition. **(I)** MitoPathways GSEA of patient LCL proteomics data in H. **(J)** Immunoblotting of IMS-localized proteins in patient LCLs with a heterozygous FAM136A-Q76* mutation overexpressing GFP-FLAG or FAM136A-FLAG. **(K)** FAM136A stabilizes a subset of IMS proteins, including CLPB and its substrate HAX1, ENDOG, and NNT, and supports OXPHOS. OMM: outer-mitochondrial membrane. IMM: inner-mitochondrial membrane. In A, B, E, F, H, and I, Mitochondrial central dogma (black) and OXPHOS-related (orange) pathways as defined in MitoCarta3.0 are highlighted.

We next sought to confirm the function of *FAM136A* in tissues by examining organs isolated from *Fam136a* knock-out (*Fam136a*^KO^) mice that present age-related hearing loss reminiscent of Ménière’s disease^29^. We obtained brain, heart, and liver from aged (20-26-month-old) *Fam136a*^KO^ mice and analyzed their protein composition using quantitative proteomics (**Table S4B-D**). Similarly to K562 cells, we observed a decreased abundance of selected IMS proteins, including ClpB, Hax1, and Nnt (**Figure S4A**), as well as a global decrease in OXPHOS protein abundances in the heart, but not in the brain or liver, suggesting tissue specificity (**Figure 4E-F and S4B-C**). However, unlike the observations in K562 cells, we did not observe any decrease in mitochondrial central dogma proteins in the three mouse tissues. Rather, the heart showed a modest enrichment of these proteins (**Figure 4E-F**), suggesting a compensatory mechanism. Supporting a role for *Fam136a* in mitochondrial function *in vivo*, we also observed a significant upregulation of proteins involved in the integrated stress response (ISR) in both heart and liver, including in Phgdh, Asns, and Mthfd1l^30–33^ (**Figure 4G and S4D**), a typical systemic response to mitochondrial dysfunction^34–36^.

Finally, we tested the relevance of our findings in patient cells. We obtained a lymphoblastoid cell line (LCL) isolated from a Ménière’s disease patient^23^ heterozygous for the FAM136A-Q76* mutation (**Figure S4E**). To test if increased levels of FAM136A could revert negative effects on IMS proteostasis, we reintroduced a wild-type copy of FAM136A-FLAG in the patient LCLs and subjected them to quantitative proteomics (**Figure 4H-I, Table S4E**). Importantly, we found that while FAM136A-FLAG reintroduction significantly increased OXPHOS proteins, it also supported HAX1, CLPB, and ENDOG protein abundance in patient cells (**Figure 4J**). Taken together, and confirming results from our genetic screen, we conclude that the depletion of *FAM136A*/*Fam136a* interferes with IMS proteostasis and causes OXPHOS defects in cell lines, mice, and patients, with tissue-specific variability reminiscent of MDs (**Figure 4K**).

## Discussion

We have presented an updated inventory of human genes required for OXPHOS. Our inventory combines data obtained from our previous death^8^ and new growth screens, two distinct screening methodologies that we show are potent and complementary at identifying MD genes. Genome-wide CRISPR-Cas9 screening has emerged as an extraordinary tool for functional genomics, and to date, several screening modalities have been developed, based on cell viability, microscopy, or flow cytometry-assisted sorting^37^, and those screening modalities have been used in combination with environmental conditions relevant to mitochondria, such as low glucose^38,39^, human plasma-like medium^40^, uridine as a carbon source^14^, vitamins^41^, or varying oxygen levels^42^. Here, we directly compared galactose-sensitized death and growth screening and benchmarked their efficiency using a curated list of 307 protein-coding nuclear-encoded MD genes^4^ and MitoPathways^19^. We found that while both screening modalities perform equally in identifying MD genes and some major mitochondrial pathways, they show differences in their ability to retrieve others. The main factor explaining these differences is the duration of the screens, now performed over 21 days for growth screening (> 10 cell divisions), as opposed to 24 hours for death screening (**Figure 1A**). We noticed that death screening was more potent at identifying central dogma genes, raising the possibility that cells might adapt to mild mtDNA defects over time. Conversely, we identified several new pathways, such as those related to mitochondrial dynamics and protein homeostasis, only identified with growth screening. A possible explanation resides in the fact that depletion of those genes leads to a progressive loss in mtDNA or to the accumulation of unprocessed or misfolded protein products, which would present a phenotype only with time, reminiscent of late-onset MDs. Our updated inventory identified 157 of the 307 protein-coding genes known to cause MDs, and it remains to be clarified why a significant portion of MD-associated genes were not detected by either screen. Possible reasons include cell line specificity, metabolic complementation from supplements included in the screen medium, functional redundancy that would mask phenotypes in our single gene targeting approach, or genes involved in secondary MDs. Our comparison of growth and death screening highlights the strong complementarity of these methods. Using both strategies in parallel could generalize to other phenotypes and better support gene identification in the future.

Here, we have focused on *FAM136A*, an uncharacterized MitoCarta3.0 gene involved in a presumed non-mitochondrial disorder. Previous work has shown that FAM136A localizes in the IMS in proximity to several IMS-exposed proteins^24^, and while a *Fam136a*^KO^ mouse model has been generated and presents with age-related hearing loss reminiscent of Ménière’s disease^29^, the role of this gene in mitochondria was not known. We have shown here that the depletion of *FAM136A* results in the depletion of several IMS proteins across all our models, with among them HAX1 and CLPB, two MD-associated genes^43,44^, indicating a primary role in IMS protein homeostasis. *FAM136A* depletion also resulted in OXPHOS protein loss in human cultured K562 cells, in patient-derived LCLs, as well as in the heart of *Fam136A*^KO^ mice, where it is accompanied by MD-like ISR signaling, but not in the liver or the brain. This variability is a common feature of MDs, likely caused by organ-to-organ variation in metabolic sensitivity and the existence of compensatory mechanisms^36^.

A *FAM136A* nonsense mutation is associated with a dominant form of fMD, and based on our work, we propose that mitochondria may play a role in the etiology of the disease. While our K562 and mouse models are based on a homozygous genetic inactivation, the *FAM136A* mutation seen in patients is heterozygous, and the associated disease presents with an autosomal-dominant inheritance pattern, with symptoms presumably restricted to the inner ear. Based on our observations, we propose that heterozygous mutations in *FAM136A* represent a mild presentation of an MD and that patients carrying homozygous inactivating mutations in *FAM136A* would present with severe, multi-organ MD.

Our work highlights the power of nutrient-sensitized screening to discover human disease genes. While whole genome sequencing on patients presenting with a suspicion of MD currently exhibits only a limited success rate, we believe our catalog of genes will be an important resource for geneticists and clinicians, allowing them to prioritize variants based on our experimental evidence, as well as for mitochondrial researchers.

## Acknowledgments

The authors would like to thank V. Mootha, T. Roger, M. Quadroni, and members of the Jourdain lab. This work was supported by grants from the Swiss National Science Foundation (310030_200796 to AAJ), the Fondation Suisse de Recherche sur les Maladies Musculaires (FSRMM, to AAJ), the Novartis foundation for medical-biological research (to AAJ), a grant from the American Hearing Research Foundation (to AL) and two EMBO postdoctoral fellowships (ALTF 286-2022 to ML and ALTF 944-2022 to ER). Mass spectrometry-based proteomics work was performed by the Protein Analysis Facility of the Faculty of Biology and Medicine, University of Lausanne, Lausanne, Switzerland.

## Inventory of Supplemental Material

- **Table S1**: Galactose-sensitized CRISPR-Cas9 growth screen data, related to Figure 1
- **Table S2**: Updated inventory of human genes essential for OXPHOS data, related to Figure 2
- **Table S3**: Immunoprecipitation and mass spectrometry of FAM136A-FLAG, related to Figure 4
- **Table S4**: Quantitative proteomics data of *FAM136*^KO^ cells, organs from *Fam136*^KO^ mice, and patient LCLs, related to Figure 4
- **Key Resources Table**

## Supplemental Figures and Legends

**Figure S1:**
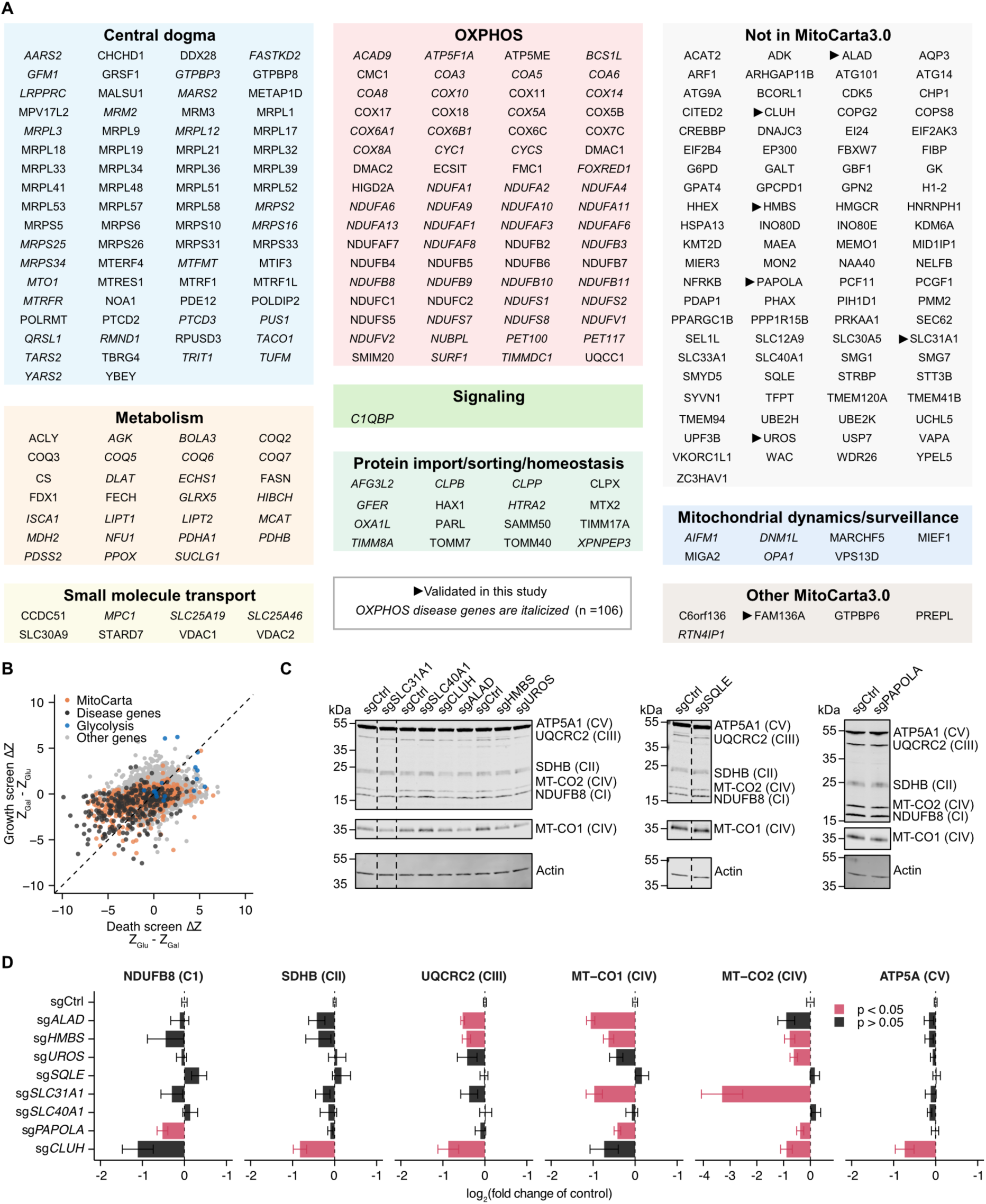
Galactose-sensitized CRISPR-Cas9 growth screen inventory and validation, related to Figure 1. **(A)** 291 growth screen hits (genes with Z < −2 x standard deviation), with 200 MitoCarta3.0 and 91 non-MitoCarta3.0 genes. MitoCarta3.0 genes are further divided into 7 high-level MitoPathways^19^. Italics: 106 genes involved in MDs. Arrowheads: genes investigated in this study. HAX1 and GTPBP8 are not represented in MitoCarta3.0 and were manually assigned based on previous work^45,46^. **(B)** Gene representation of the genome-wide CRISPR-Cas9 death vs growth screen showing DZ-scores. N = 10,775 genes represented in both datasets **(C)** Representative immunoblot showing 6 protein subunits of the mitochondrial respiratory chain in the indicated cell lines. Each panel represents an individual immunoblot, and vertical dotted lines indicate line splicing. **(D)** Immunoblot quantification (normalized to actin) of C, from n = 4-8 transduction replicates. Bars represent mean log2 fold-changes compared to their matched controls. Error bars represent mean +/- SEM. Statistical analysis: one-sample t tests.

**Figure S2:**
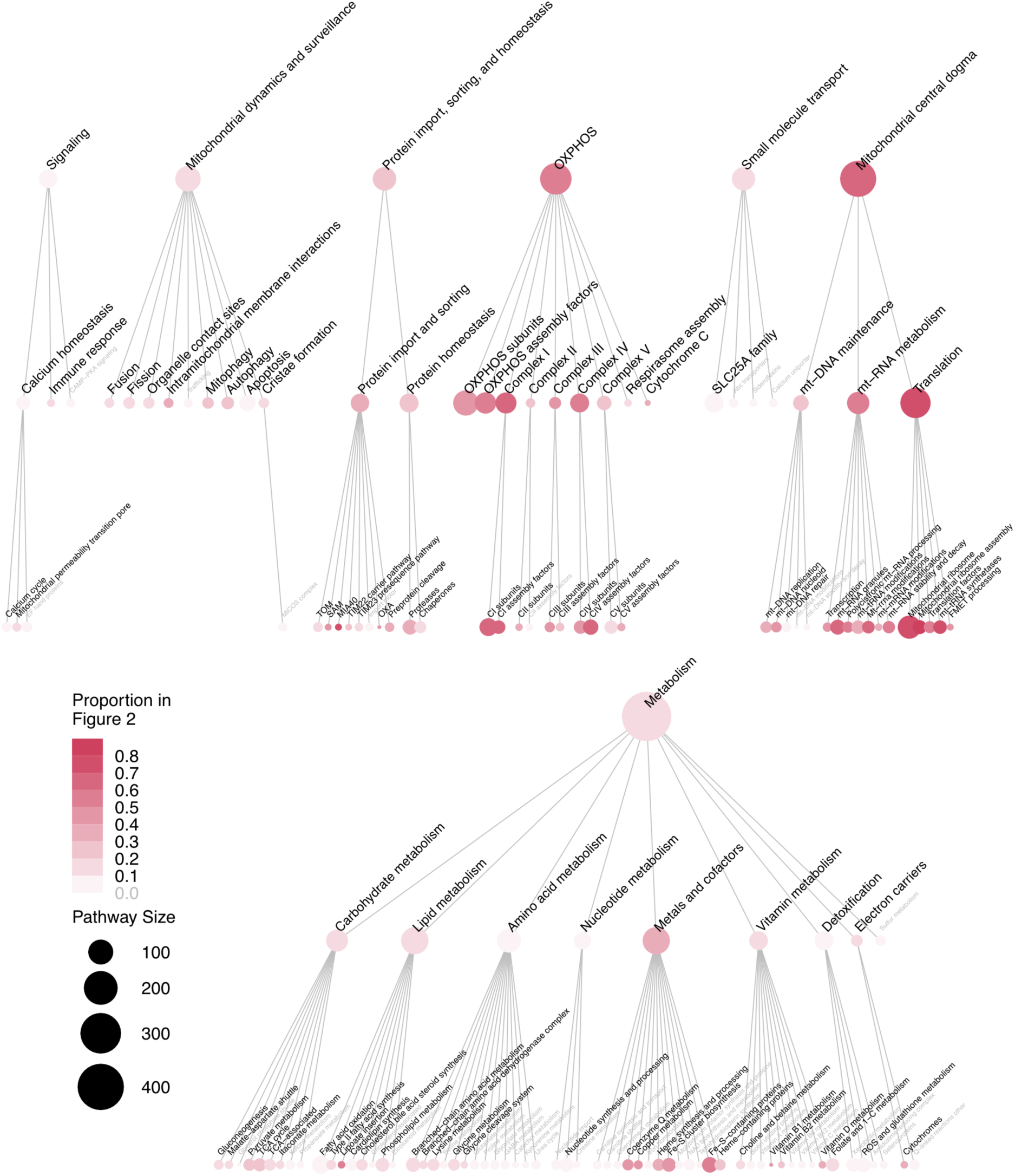
MitoPathways analysis of the updated inventory of human genes essential for OXPHOS, related to Figure 2. Circle size reflects the total size of each gene set in MitoPathways. Color gradient indicates the proportion of each gene set that is identified in the combined death and growth screens (genes listed in Figure 2). Gene sets in grey indicate sets with no identified genes in either screen.

**Figure S3:**
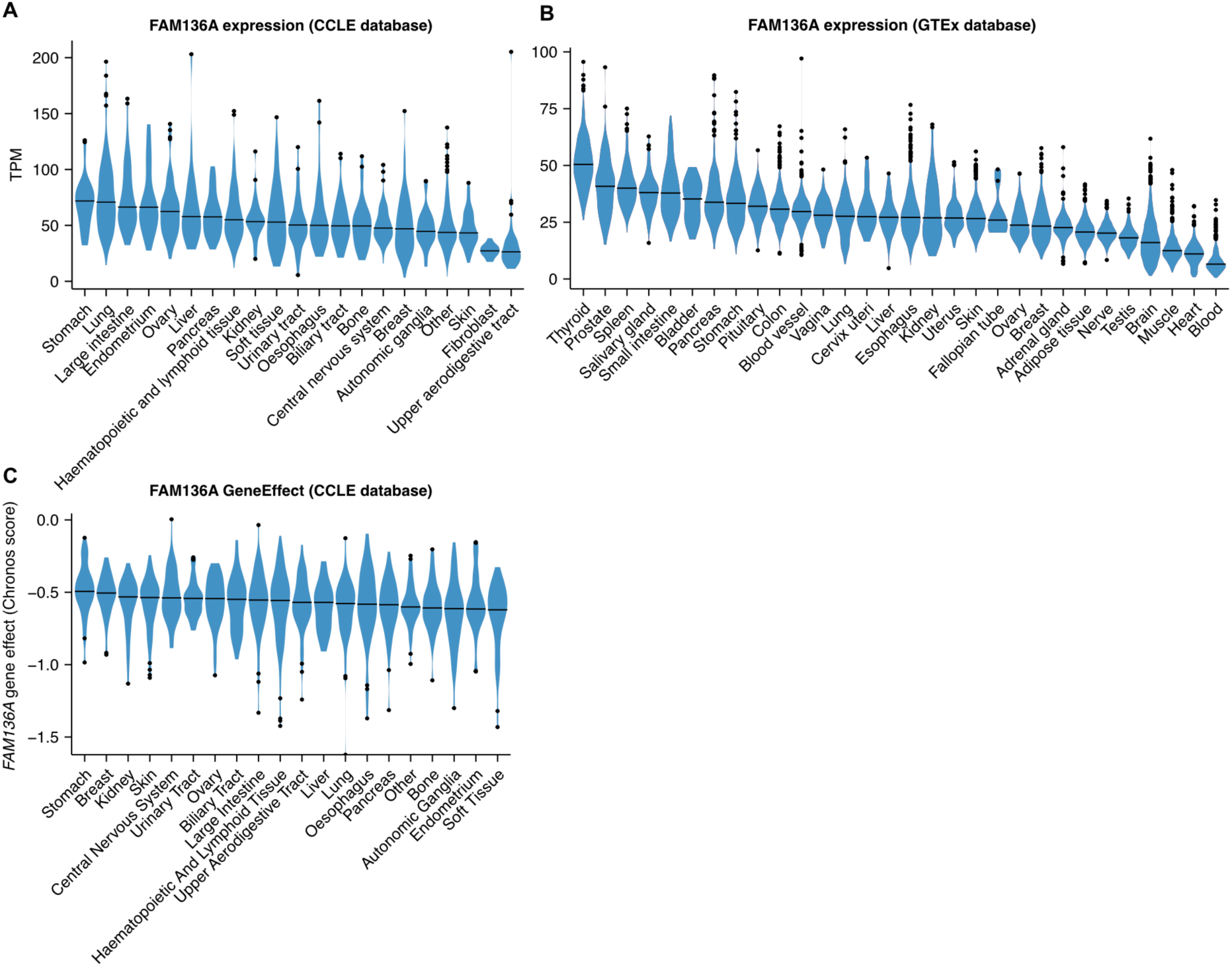
*FAM136A* expression and essentiality score across the CCLE and GTEx databases, related to Figure 3. **(A)** Violin plot showing CCLE *FAM136A* expression by tissue of origin. **(B)** Violin plot showing GTEx *FAM136A* expression by tissue. **(C)** Violin plot showing CCLE *FAM136A* expression by tissue of origin. Tissues with less than 20 representative cell lines are categorized as “Other”. Horizontal lines indicate medians, and points represent outlier values (outside 1.5 times the interquartile range above the 0.75 quantile or below the 0.25 quantile). TPM: transcripts per million.

**Figure S4:**
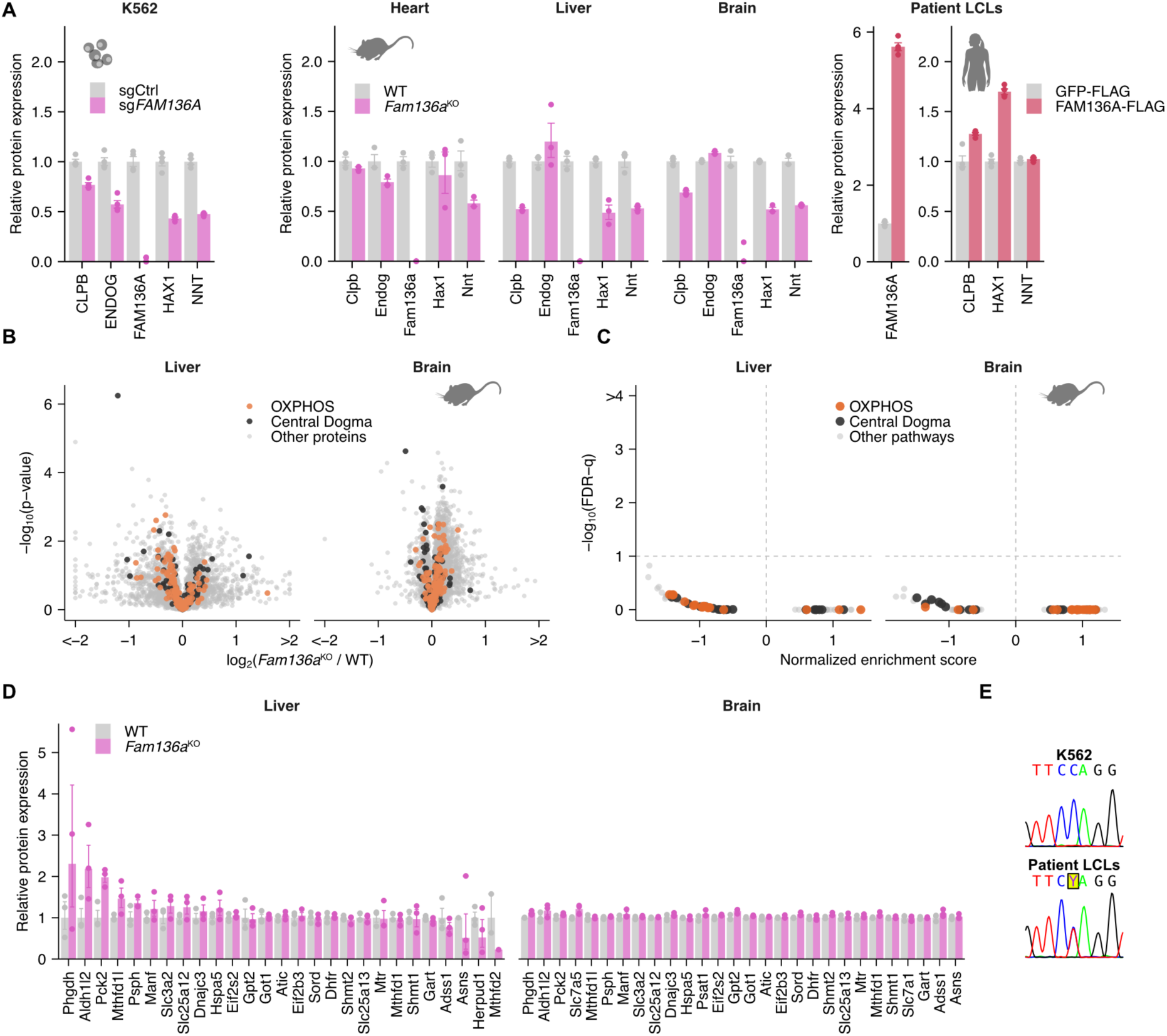
Additional proteomics of analysis of *FAM136*^KO^ cells, organs from *Fam136*^KO^ mice, and patient LCLs, related to figure 4. **(A)** Bar plots of selected mitochondrial intermembrane space-localized proteins. n = 4 independent infections (K562), 3 mice per condition (mouse tissues), or 4 replicates (LCLs)**. (B)** Quantitative proteomics comparing protein levels in liver and brain tissues of *Fam136a*^KO^ mice vs controls (WT). Proteins involved in mitochondrial central dogma (black) and OXPHOS (orange) as defined in MitoCarta3.0 are highlighted. n = 3 mice per condition. **(C)** MitoPathways GSEA of mouse tissue proteomics data in B. **(D)** Bar plots showing log_2_(fold changes) of proteins involved in the ISR in the mouse liver and brain tissue proteomics datasets in B. ISR genes were manually assigned based on previous studies^30–33^ (**Supplementary table S4F**). **(E)** Sanger sequencing of the mutated locus in patient LCLs compared to K562 cell lines. Y indicates a pyrimidine (heterozygous C-T mutation).

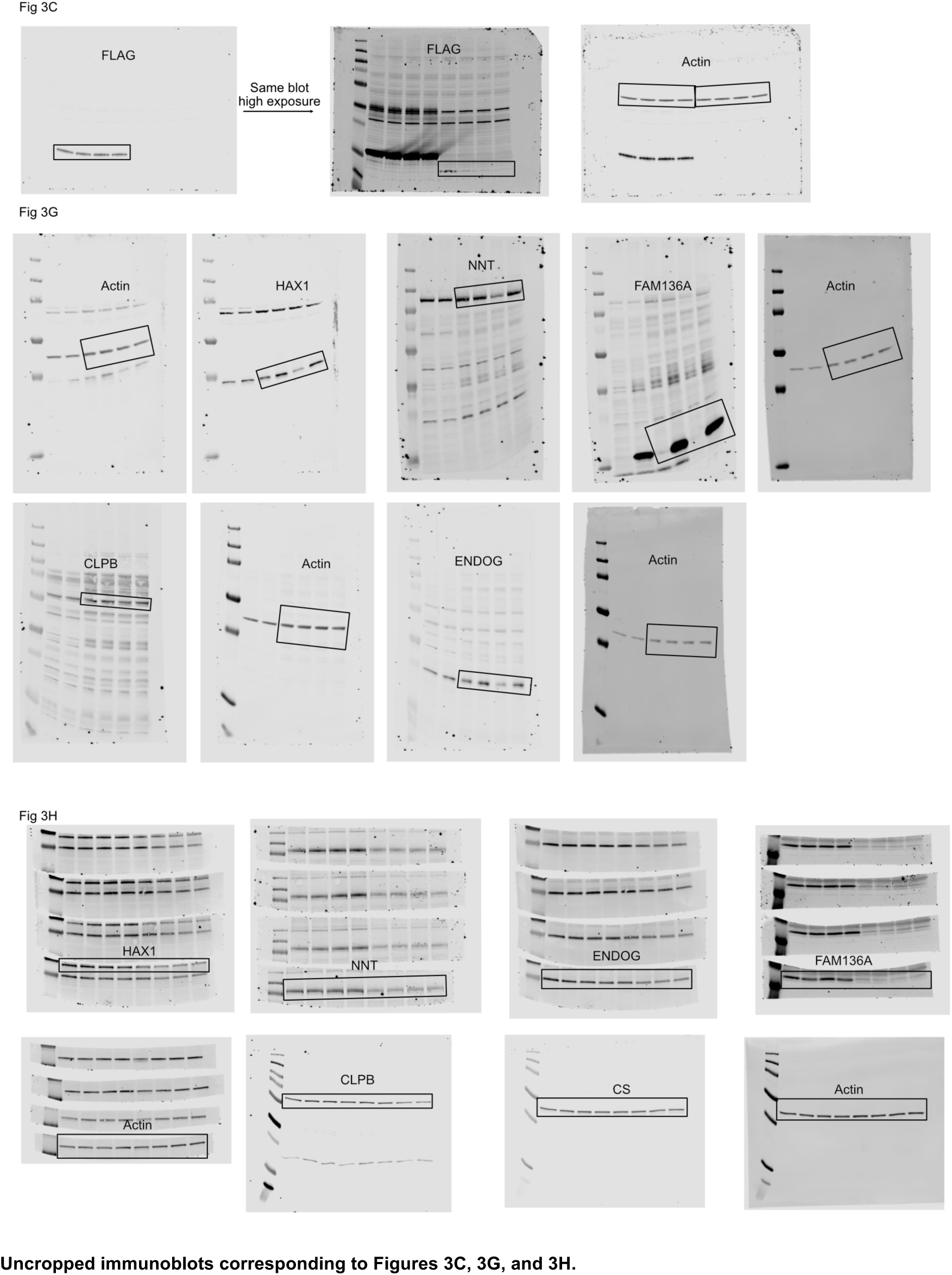
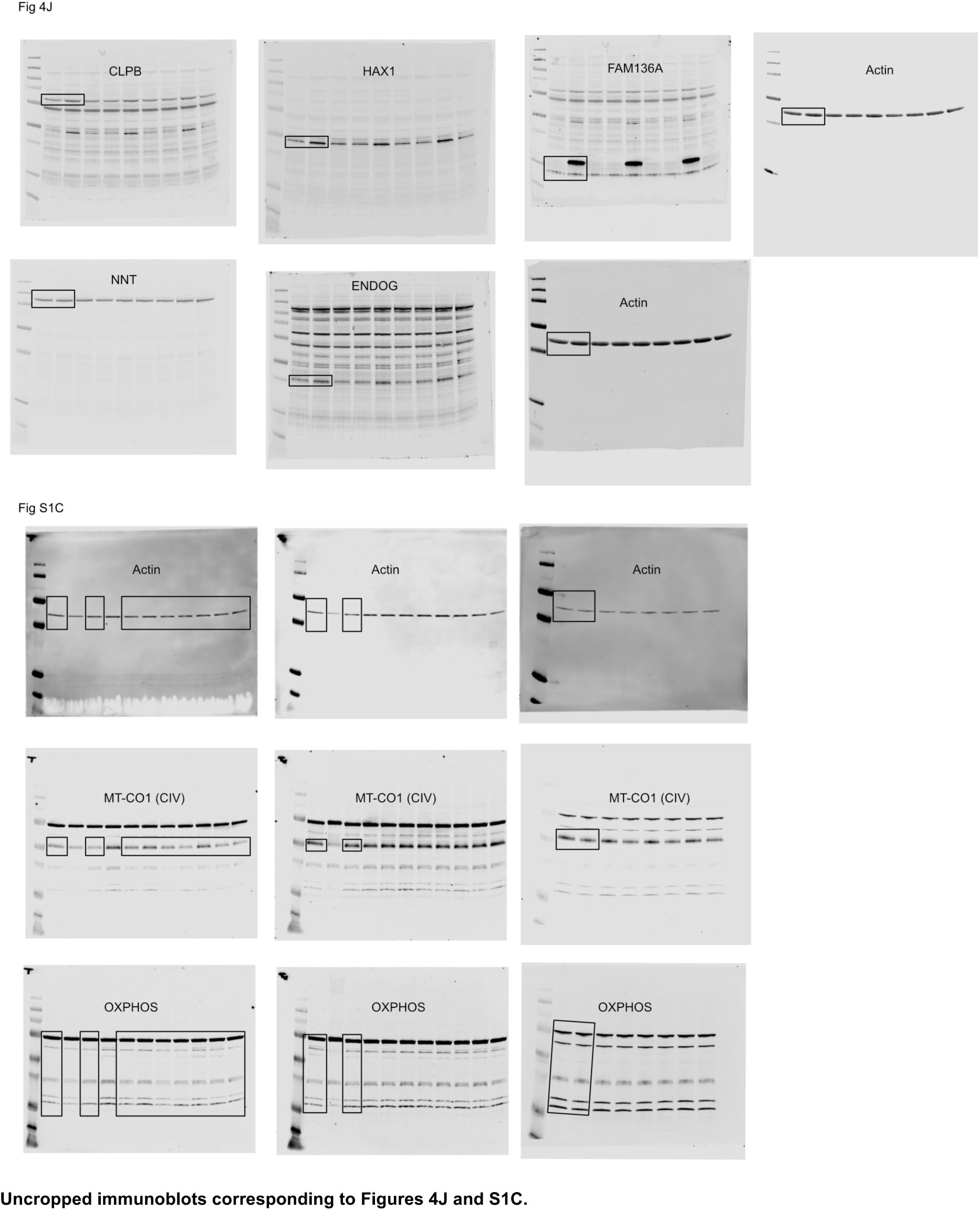

## Author contributions

**MH:** investigation (lead); methodology (equal); validation (lead); data curation (equal); project administration (equal); formal analysis (supporting); visualization (supporting); writing (original draft preparation, equal); **MMF:** investigation (supporting); methodology (equal); validation (supporting); data curation (equal); project administration (supporting); formal analysis (lead); visualization (lead); writing (original draft preparation, equal); **ER:** investigation (supporting); writing (reviewing and editing, equal); funding acquisition (supporting); **ML:** investigation (supporting); writing (reviewing and editing, equal); funding acquisition (supporting); **SC:** investigation (supporting); writing (reviewing and editing, equal); **JCL:** investigation (supporting); writing (reviewing and editing, equal); **NL:** investigation (supporting); **SM:** supervision (supporting); writing (reviewing and editing, equal); **JALE:** supervision (supporting); writing (reviewing and editing, equal); **AL:** supervision (supporting); writing (reviewing and editing, equal); **AAJ:** conceptualization (lead); investigation (supporting); methodology (equal); project administration (equal); supervision (lead); funding acquisition (lead); writing (original draft preparation, equal).

## Declaration of generative AI and AI-assisted technologies

No generative AI or AI-assisted technology was employed for conceptualization or investigation in the course of this project.

## Data availability

The analyzed mass spectrometry proteomics data are available in the supplemental tables and the raw data have been deposited to the ProteomeXchange Consortium via the PRIDE^47^ partner repository with the dataset identifiers PXD060163 (K562), PXD060280 (IP-MS), PXD060278 (mouse tissue), and PXD060152 (patient LCL).

## Methods

### Cell lines

K562 (CCL-243; ATCC), 293T (CRL-3216; ATCC), U2-OS (HTB-96, ATCC). fMD patient LCLs were from^23^. Cells were regularly tested for mycoplasma and the genotype of fMD LCLs was confirmed by Sanger sequencing (below).

### Cell Culture

Cells were maintained in DMEM high glucose, GlutaMAX (Gibco, 31966-021) (for K562, 293T), McCoy’s 5A (Gibco, 26600023) (for U2-OS), or RPMI 1640 (Gibco, 61870-010) (for LCL) medium supplemented with 10% fetal bovine serum (FBS, Thermo Fisher Scientific, 17479633) and 100 U/mL penicillin/streptomycin (BioConcept, 4-01F00-H) under 5% CO2 at 37°C. Cells were counted using a Vi-CELL BLU Instrument cell counter (Beckman Coulter, C19196), and only viable cells were considered.

### CRISPR-Cas9 screen

Genome-wide CRISPR-Cas9 screening was performed in duplicate using K562 cells and a lentiviral-carried Brunello library (Genome Perturbation Platform, Broad Institute)^48^ containing 76,441 sgRNAs, as described before^14^. Cells were infected with a multiplicity of infection of 0.3 and at 500 cells per sgRNA in the presence of 10 µg/mL polybrene (Sigma-Aldrich, TR-1003-G). After 24 h, cells were transferred to a culture medium containing 2 µg/mL puromycin (InvivoGen, ant-pr-1) and incubated for an additional 48 h before removing puromycin. On day 10, the cells were plated in glucose-free DMEM (Gibco, A1443001) containing 10% dialyzed FBS (Sigma-Aldrich, F0392), 50 µg/mL uridine (Sigma-Aldrich, U3003), 1 mM pyruvate (Thermo Fisher Scientific, 11360070), and 100 U/mL penicillin-streptomycin, and supplemented with 25 mM of either glucose (Sigma-Aldrich, G7021) or galactose (Sigma-Aldrich, G5388) at a concentration of 10^5^ cells per mL and with 1,000 cells per sgRNA. Cells were passaged every 3 days for 3 weeks and, on day 31, 1,000 cells per sgRNA were harvested.

For DNA isolation, total genomic DNA was isolated from cells using a NucleoSpin Blood kit (Machery Nagel, 740954.20) using the manufacturer’s recommendations. Barcode sequencing, mapping and read count were performed by the Genome Perturbation Platform (Broad Institute).

Data were analyzed using Z-score^49^. Briefly, log2-transformed reads per million (log2(sgRNA-reads / total reads x 10^6^ + 1) were calculated for each sgRNA and normalized to the pre-swap controls (day 7 after library infection). Log2(fold changes) were then averaged per gene and condition (galactose or complete medium). For each condition, null distribution parameters were calculated based on the distribution of genes of low expression (TPM < 1 in the CCLE 24Q2 expression dataset^25^, and log2(fold changes) were Z-normalized using the mean and standard deviations of the null distributions. Only genes with TPM > 1 in the CCLE 24Q2 expression dataset are reported here.

### Updated inventory of human genes required for OXPHOS

300 genes from the previously published CRISPR death screen^8^ with an FDR q-value below 0.3 were combined with 291 genes from the growth screen with a ΔZ-score (Z_gal_ – Z_glu_) below −2 x standard deviation of the ΔZ-scores, yielding 481 unique genes. Genes were categorized according to the top-level MitoCarta3.0 MitoPathways terms (Mitochondrial central dogma; Metabolism; OXPHOS; Small molecule transport, Protein import, sorting, and homeostasis, Mitochondrial dynamics and surveillance; or Signaling). 43 of the 481 genes were annotated with more than one top-level MitoPathway term and for these, one pathway was manually selected based on previous work. Confidence scores for each gene were calculated as their ΔZ rank sums i.e., rank (death screen) + rank (growth screen).

### CCLE dependency correlations and GSEA

Chronos gene effect scores (24Q2) were downloaded from the Cancer Cell Line Encyclopedia (CCLE) Dependency Portal (DepMap)^50^. *FAM136A* Chronos scores were correlated (Pearson correlation coefficient) with Chronos scores of all other CCLE genes, excluding genes with Chonos scores in less than 25 of the 1,150 CCLE cell lines. Correlation coefficients were subsequently exported for gene set enrichment analysis.

### GTEx and CCLE gene expression analysis

Gene TPMs were downloaded from the GTEx portal (GTEx Analysis V8, GTEx_Analysis_2017-06-05_v8_RNASeQCv1.1.9_gene_tpm.gct.gz, accessed 30.9.2024) and the CCLE DepMap portal^25^ (OmicsExpressionProteinCodingGenesTPMLogp1 24Q2, accessed 11.10.2024).

### Cloning of sgRNAs to *FAM136A* for gene-specific CRISPR-Cas9 knockouts

The two best *FAM136A* sgRNAs (ACCGCATGGTGCACCGGGCCAGG, ACCGCTGGTGCACCTGCTTCATGG) from the Brunello library were ordered as complementary oligonucleotides (Integrated DNA Technologies) and cloned in pLentiCRISPR v2^51^. sgRNAs targeting OR11A1 (CACCGGTGATGCCAAAAATGCTGGA) and OR2M4 (CACCGCCATAAGGGACAGTTGACTG) (two unexpressed genes in K562 cells) were used as a negative control.

### Cloning of human wild-type and mutant FAM136A-FLAG

FAM136A-3xFLAG and FAM136A_1-75_-3xFLAG cDNAs were ordered as gBlocks (Integrated DNA Technologies) and were amplified with the primers ATCGATCGGGATCCATGGCA and CGATCGATGCGGCCGCTCAC (Integrated DNA Technologies). PCR products were purified (Qiagen, 28706) and digested with BamHI-HF and NotI-HI (NEBNew England Biolabs, R3136S and R3189S). Both inserts were cloned into pLV-EF1a-IRES-Hygro plasmid (Addgene, 85134).

### Lentiviral transduction

Lentiviruses were produced from pLentiCRISPR v2 (Addgene, 52961)^51^ and pLV-EF1a-IRES-Hygro (Addgene, 85134)^52^ according to Addgene’s protocol, and 24 to 48 h post-infection, cells were selected with 2 µg/mL puromycin (InvivoGen, ant-pr-1) or 200 µg/mL hygromycin B (Sigma-Aldrich, 3274) for 2 to 7 days. Cells were then maintained in routine culture medium for 15 days post-transduction before analysis. Gene disruption efficiency was verified by immunoblotting.

### Immunoblotting and quantification

Cells were harvested, washed twice in PBS, and lysed for 10-20 min on ice in RIPA buffer (25 mM Tris pH 7.5, 150 mM NaCl, 0.1% SDS, 0.1% sodium deoxycholate, 1% NP40 analog) supplemented with Protease Inhibitor Cocktail (Sigma-Aldrich, PIC0002) and Universal Nuclease (Life Technologies, 88702). Lysates were spun down at 1400 *g* for 10 min at 4 °C, and supernatants were moved to new 1.5 mL Eppendorf tubes. Protein concentration was determined from total cell lysates using the DC protein assay (Bio-Rad, 5000111). Lysates were complemented with 5x SDS-PAGE loading buffer (4% SDS, 250mM Tris pH 6.8, 0.6 mg/mL bromophenol blue, 2.3 M ß-mercaptoethanol, 33.3% glycerol enhydrous) and gel electrophoresis was done on Novex™ 10-20%, Tris-Glycine Plus WedgeWell™ gels (Invitrogen, XP10200BOX) before transfer by wet transfer on nitrocellulose membranes (Sigma-Aldrich, GE10600004). Equal loading was confirmed by Ponceau staining (Thermo Fisher Scientific, A40000278). Blocking and antibody incubation were performed in 5% milk powder TBS supplemented with 0.1% Tween® 20 (PanReac AppliChem, A4974,0500). Primary antibodies were diluted 1:100-1:5000 and secondary fluorescent-coupled secondary antibodies were diluted 1:10,000 in the same buffer. Washes were done in TBS supplemented with 0.1% Tween® 20. Imaging and quantification were done using Image Studio version 4.0 on an Odyssey CLx analyzer (LI-COR®).

### Protein stability assays

K562 or LCL cells were incubated for 0 to 6 hours in a medium containing 0.05 mg/mL cycloheximide (Sigma-Aldrich, 01810). After the desired time, cells were harvested by centrifugation, PBS-washed twice, and lyzed for immunoblotting as described above.

### Sanger sequencing

Genomic DNA was isolated from LCLs using QIAamp DNA Mini Kit (Qiagen, 51304). The region of the *FAM136A* gene containing the expected mutation was then amplified by PCR using primers GGACTACAGTGCTCTGTCTAGG and AGCTGTTGTGAGGACAGCCA. The PCR product was purified using QIAquick® Gel Extraction Kit (Qiagen, 28706) and further sequenced (Sanger sequencing) using the same primers. Sequencing results were visualized using the sangerseqR package^53^ in R version 4.4.0^54^.

### Oxygen consumption assays

One XFE96/XF Pro Sensor Cartridge from the Agilent Seahorse XFe96/XF Pro FluxPak (Agilent, 103792-100) was pre-hydrated using 200 µl/well Agilent Seahorse XF Calibrant (Agilent, 100840-000) and XF Hydrobooster, then incubated overnight at 37 °C without CO_2_. One hour before the Seahorse assay, 125,000 K562 or LCL cells/well were seeded in 175 µl of Agilent Seahorse XF DMEM Medium (Agilent, 103575-100) or Agilent Seahorse XF RPMI Medium (Agilent, 103576-100), respectively, with 25 mM D-(+)-glucose and 2 mM stable glutamine 100x (L-Ala-L-Gln, 200mM) (BioConcept, 5-10K50-H), and incubated at 37 °C without CO_2_ for one hour. 25 µl of oligomycin A (Tocris, 4110), carbonyl cyanide m-chlorophenylhydrazone (Sigma-Aldrich, C2759), and antimycin A (Sigma-Aldrich, A8674) were loaded into pores A, B, and C of the sensor cartridge for final in-well concentrations of 2 µM, 1.5 µM, and 1.6 µM, respectively. Pore D was left empty. Then, the sensor cartridge was loaded into an Agilent Seahorse XFe96 Analyzer for calibration. After calibration, the XFe96/XF Pro Cell Culture Microplate (Agilent, 103794-100) was loaded into the XFe96 Analyzer, and the Mito Stress Test was conducted and analyzed using Seahorse Wave Controller 2.6.3.5 software (Agilent).

### Protein structure predictions

FAM136A and FAM136A_1-75_ structures were predicted using AlphaFold 3^55^ and visualized using UCSF Chimera version 1.18^56^.

### Quantitative cell proteomics

K562 or LCL cells were harvested, washed twice in PBS, and pellets were frozen at −80°C. Cell pellets (5×10^6^ cells) were lysed in miST lysis buffer (1% Sodium deoxycholate, 10 mM DTT, 100 mM Tris pH 8.6) and heated for 10 min at 75 °C. Sample preparation, peptide fractionation, liquid chromatography (LC)-MS, and data processing were carried out essentially as reported previously^57^. All LC-MS analyses were done using a TIMS-TOF Pro (Bruker, Bremen, Germany) mass spectrometer coupled to an Ultimate 3000 RSLCnano HPLC system (Dionex) or an EvoSep One LC system (EvoSep, Odense, Denmark).

Raw MS data were processed with Spectronaut version 18.4 or 19.4 (Biognosys, Schlieren, Switzerland). A peptide library was constructed from data-dependent LC-MS data from pooled sample aliquots based on the Swiss-Prot human proteome (20,375 sequences, retrieved January 7^th^, 2022, for K562; 82’499 sequences, retrieved February 4^th^, 2024, for LCLs). The library contained 126,853/145,459 (for K562 or LCLs, respectively) precursors mapping to 92,188/81,168 stripped sequences, of which 88,134/37,290 were proteotypic. These corresponded to 8,004/8,335 protein groups (8,096/12,584 proteins). Of these, 774/952 were single hits (one peptide precursor). In total 749,333/869,060 fragments were used for quantitation. Peptide-centric analysis of data independent LC-MS data resulted in a total of 120,266/138,375 precursors, mapping to 7,838/8,101 protein groups; 101,203/115,091 precursors (7,231/7,570 protein groups) were quantified in all samples. Downstream analyses were performed using an in-house-developed tool (https://github.com/UNIL-PAF/taram-backend). Contaminant proteins were removed, and protein group abundances were log2-transformed. Only proteins quantified in all samples in one group were kept. Missing values were imputed based on a normal distribution with a width of 0.3 standard deviations (SD), downshifted by 1.8 standard deviations relative to the median. *p*-values were calculated by Student’s t-tests and corrected using the Benjamini and Hochberg method. Data were visualized using the ggplot2 package^58^ in R version 4.4.0^54^.

### Animals

*Fam136a*^KO^ mice (Fam136a^tm1a(KOMP)Wtsi^, Strain of origin: C57BL/6N-A^tm1Brd^) were obtained from the Wellcome Trust Sanger Institute (Hinxton, UK)^59^. Mice were placed in cages containing up to five mice and fed a regular diet of mouse chow and water. After breeding, tail snips of the mice were obtained at weaning (postnatal day 21, PD21) and DNA-sequenced (Transnetyx, Inc., Cordova, TN) to determine genotype. Ear punches were used to uniquely identify individual mice in each cage. Organ isolation was performed from wild-type (two female and one male) and homozygous *Fam136a*^KO^ mice (two female and one male) aged 20-26 months old. All animal experiments were carried out in the Department of Anatomy and Cell Biology of University Illinois Chicago. Animal experiments were approved by the University of Illinois at Chicago IACUC review board (Protocol #20-103) and conformed to the Guide for the Care and Use of Laboratory Animals.

### Quantitative mouse tissue proteomics

Mice were anesthetized and transcardiacally perfused with PBS. Tissues were excised and flash-frozen in liquid nitrogen. Tissue lysis and proteomics analysis were performed at the Protein Analysis Facility of the Faculty of Biology and Medicine, University of Lausanne, Switzerland. Frozen tissues were disrupted by agitation with ceramic beads in an excess of 80% methanol prechilled at −20 °C (Fastprep system). After incubation for 1 h at −20 °C, the solvent supernatant was removed. The tissue paste was resuspended in cold miST buffer and shaken again 3×30 s in the Fastprep with 2 min on ice between steps. The resulting supernatant was transferred to new tubes and heated at 75 °C for 10 min. Extracts were diluted 1:1 with miST buffer and vortexed 2 min before protein concentration quantitation.

Sample digestion was performed according to a modified version of the iST method^60^, as previously described^57^. For spectral library generation (only heart and liver samples), aliquots from all samples were pooled and fractionated into six fractions by off-line basic reversed-phase (bRP) using the Pierce High pH Reversed-Phase Peptide Fractionation Kit (Thermo Fisher Scientific, 84868). The 6 fractions were collected in 7.5%, 10%, 12.5%, 15%, 17.5%, and 50% acetonitrile with 0.1% triethylamine (∼pH 10). Dried bRP fractions were reconstituted in 2% acetonitrile with 0.5% trifluoroacetic Acid (TFA) and injected for LC-MS/MS analyses. LC-MS/MS analyses were performed using a timsTOF Pro (Bruker, Bremen, Germany) mass spectrometer interfaced through a nanospray ion source to an EvoSep One LC system (EvoSep, Odense, Denmark). Peptides were separated on a reversed-phase C18 column (EvoSep, EV1137) at 0.22 µl/min with a gradient from 0 to 35% acetonitrile in water with 0.1% formic acid over 88 min. Data-dependent acquisition (DDA) was performed using a method similar to a standard TIMS PASEF method^61^ with ion accumulation for 100 ms. The duty cycle was kept at 100%. Precursor ions in the ion mobility range from 1/k0 = 0.8 to 1.3 and between *m/z* = 400 to 1200 were selected. Up to 10 precursors were targeted per TIMS scan. Precursor isolation windows were set to 2 or 3 *m/z,* below or above m/z 800, respectively. The threshold for minimum intensity for precursor selection was set to 2,500. When the inclusion list allowed, precursors were targeted more than once to reach a minimum total intensity of 20,000. Collision energy was linearly ramped based on the 1/k0 values from 20 (at 1/k0=0.6) to 59 eV (at 1/k0 = 1.6). The total scan cycle duration including one survey and 10 MS2 TIMS scans was 1.16 s. Retargeting of precursors was allowed when their signal increased by at least a factor of 4. Upon being selected, precursors were excluded for 60 s. The resolution was approximately 35,000. For liver, heart, and brain samples, data-independent acquisitions (DIA) were performed as previously reported^62^, with the same instrument parameter changes as the DDA method described above. Per cycle, the mass range 400-1,200 m/z was covered by a total of 32 windows, each 25 Th wide and a 1/k0 range of 0.3. Collision energy and resolution settings were the same as in the DDA method. Two windows were acquired per TIMS scan (100 ms) so that the total cycle time was 1.8 s.

Data processing was performed separately for the brain samples or the heart and liver samples. Identification of peptides in brain samples directly from DIA data was performed with Spectronaut 18.6 with the Pulsar engine using the “deep” setting and searching the reference mouse proteome (www.uniprot.org, 54,822 sequences, accessed February 14^th^, 2024). For identification, tryptic peptides comprised of 7-52 amino acids, with a maximum of 2 missed cleavages were considered. Cysteine carbamidomethylation was set as a fixed modification and methionine oxidation and protein N-terminal acetylation as variable modifications. Identified peptides and protein groups were filtered at a false discovery rate of 1%. Peptide ion mobility was predicted using a deep neural network and used in scoring. The generated library contained 155,908 precursors. A peptide-centric analysis of the data was performed with Spectronaut 18.6 using the generated library. Peptide quantitation was based on the extracted ion chromatogram area. Between 1 and 3 precursors, from which the median value was calculated, were considered for each peptide. Protein group quantifications were derived from inter-run peptide ratios based on the MaxLFQ algorithm^63^. Global normalization was based on peptide medians. Overall, 154,960 precursors mapped to 8,882 protein groups were quantified in the brain tissue. 135,733 precursors (8,593 protein groups) were quantified in all samples. The average number of data points per peak was 8.5. For heart and liver tissue, raw data were processed with Spectronaut 18.7 (Biognosys, Schlieren, Switzerland). A library was constructed from the DDA bRP fraction data by searching the reference mouse proteome. For identification, tryptic peptides comprised of 7-52 amino acids, with a maximum of 2 missed cleavages were considered. Cysteine carbamidomethylation was set as fixed modification and methionine oxidation and protein N-terminal acetylation as variable modifications. The Pulsar engine was used for peptide identification, and protein inference was performed with the IDPicker algorithm. Identified spectra, peptides, and proteins were filtered at a false discovery rate of 1% using a decoy database. Fragments corresponding to less than 3 amino acids, fragments outside the m/z range of 300-1800, and precursors with less than 3 fragments were removed from the library, and only fragments with a minimum base peak intensity of 5% were included. Additionally, only the 6 best fragments per precursor were kept. Shared (non-proteotypic) peptides were kept in the library. The library contained 73,142 precursors mapping to 55,855 stripped sequences, of which 24,573 were proteotypic. These corresponded to 6,160 protein groups (9,205 proteins). Of these, 1,049 were single hits (one peptide precursor). In total 436,201 fragments were used for quantification. Peptide-centric analysis of heart and liver DIA data was done using Spectronaut 18.7 using the generated library. Peptide quantification of liver and heart samples was carried out similarly to the brain samples. 68,446 precursors were quantified in the heart and liver tissue dataset, and mapped to 5,994 protein groups (8,956 proteins). 34,471 precursors (3,917 protein groups) were quantified in all samples. Downstream analyses were performed as described for the quantitative cell proteomics.

### Immunoprecipitation and mass spectrometry

Immunoprecipitation was performed as previously described^64–66^. GFP-3xFLAG and FAM136A-3xFLAG were stably expressed in K562 cells and a crude mitochondrial fraction^67^ was prepared from an equal number of cells from each condition in duplicates and lysed in IP lysis buffer (50 mM Tris/HCl (pH 7.5), 150 mM NaCl, 1 mM MgCl2, 1% NP-40, 1x protease (Cell Signaling)). Lysates were cleared by centrifugation at 20,000 g for 20 min and the supernatants were saved. Washed anti-FLAG M2 magnetic beads (Sigma-Aldrich, M8823) were added to the lysate and incubated overnight at 4 °C. Beads were recovered after extensive washing, and the protein complexes were eluted with 100 μg/mL 3xFLAG peptide (Sigma-Aldrich, SAE0194) following TCA precipitation. Pellets were resuspended in SDS-PAGE loading buffer and analyzed by MS.

Protein digestion was carried out according to the SP3 protocol^68^. Briefly, protein alkylation was performed in SDS-PAGE buffer in the dark at room temperature (RT) with 32 mM iodoacetamide for 45 min. 10:1 (w:w) Sera-Mag Speed-beads (Cytiva, 45152105050250) were added, and proteins on beads were precipitated with 60% ethanol and washed 3 times with 80% ethanol. Proteins were digested with 1 µg trypsin (Promega, V5073) in 50 µl 100 mM ammonium bicarbonate for 1 h at 37 °C, and an additional 1 µg trypsin was added for a further 1 h incubation at 37 °C. Digests were transferred to fresh tubes and supplemented with 0.5% formic acid (FA) before centrifugal evaporation. Two volumes of 1% TFA in isopropanol were added, and peptide samples were applied to a strong cation exchange plate (Waters, 186001830BA). Peptides were washed in isopropanol with 1% TFA and isopropanol with 2% acetonitrile + 0.1% FA in isopropanol and eluted in 200 µl 80:19:1 Acetonitrile:water:ammonia before drying by centrifugal evaporation.

DDA was performed using a Vanquish Neo nanoHPLC system interfaced via a nanospray Flex source to a high-resolution Orbitrap Exploris™ 480 mass spectrometer (Thermo Fisher Scientific, BRE725539). Peptides were applied to a PepMap™100 C18 trap cartridge (Thermo Fisher Scientific, 160434) before separation on a C18 custom-packed column (45 cm x 75 µm inner diameter, 1.8 µm particle size, Reprosil Pur, Dr. Maisch) with a 130 min gradient from 2% to 80% acetonitrile with 0.1% FA. Full MS survey scans were performed at a resolution of 120,000. A DDA method controlled by Xcalibur™ software (Thermo Fisher Scientific, OPTON-30965) was used that optimized the number of precursors of charge states 2+ to 5+, while maintaining a fixed scan cycle of 2 s. Fragmentation was done by higher energy collision dissociation with a normalized energy of 30% at resolution 15,000. The precursor isolation window was 1.6 *m/z*, and selected precursors were excluded for 60 s from further analysis.

Data were processed using MaxQuant 2.5.1.0^69^ employing the Andromeda search engine^70^ setting cysteine carbamidomethylation as fixed modification and methionine oxidation and protein N-terminal acetylation as variable modifications. The human reference proteome used as a sequence database for searching was based on the UniProtKB reference proteome (www.uniprot.org, accessed February 2024, 82,499 sequences) and a “contaminant” database containing typical environmental contaminants and digestion enzymes. Mass tolerance was set to 4.5 ppm after recalibration for precursors and 20 ppm for fragments. Identified peptides and proteins were filtered at a false discovery rate of 1% using a reversed-sequence decoy database. Downstream analyses were performed as described for the quantitative cell proteomics.

### Immunofluorescence

U2-OS cells were plated on 12 mm glass coverslips (Menzel, MENZCB00120RA020) 24 h before transfection with 250 ng/mL of the pLV-FAM136A-WT-3xFLAG-hygro or pLV-FAM136A_1-75_-3xFLAG-hygro plasmids, using Lipofectamine 2000 (Invitrogen, 11668019) in Opti-MEM medium (Gibco, 51985-026) following manufacturer’s instructions. Fixation was done using 3.2% paraformaldehyde (PFA) (Electron Microscope Sciences, 15714) and 0.1% glutaraldehyde (GA) (Electron Microscope Sciences, 16350) in phosphate-buffered saline (PBS), followed by washing and storage in PBS. GA autofluorescence quenching was achieved by a 7-min fresh 0.5% sodium borohydride (NaBH₄) (Sigma-Aldrich, 452882) incubation. Permeabilization and blocking in 3% bovine serum albumin (BSA) (Sigma-Aldrich, A8806-1G) and 0.1% Triton X-100 (Thermo Fisher Scientific, 85111) in PBS were done for 60 min. Primary antibodies against TOM20 (Santa Cruz, sc-17764) and FLAG (Sigma-Aldrich, F7425-2MG) were diluted 1:200 in 1% BSA and incubated overnight at 4°C, followed by two 10 minutes washes in 1% BSA in PBS. Secondary antibodies Alexa Fluor™ Plus 488 anti-mouse (A32723, Thermo Fisher Scientific) and Alexa Fluor™ 568 anti-rabbit (A-11011, Thermo Fisher Scientific) diluted 1:400 in 1% BSA were incubated at RT for 1 hour and washed twice as described. Rapid rinsing in distilled water was followed by mounting in ProLong™ Gold Antifade Mountant (Thermo Fisher Scientific, P36930), cured for 24 h in the dark at room temperature.

Instant structured illumination microscopy (iSIM) was performed on a custom-built flat-fielded microscope set-up at the Manley lab in the Swiss Federal Technology Institute of Lausanne (Ecole Polytechnique FEdErale De Lausanne, EPFL), using 488-nm and 561-nm excitation lasers, a 1.49 NA oil immersion objective (Olympus, APON60XOTIRF), an sCMOS camera (PrimeBSI Photometric, 01-PRIME-BSI-R-M-16-C), and rapid vertically scanning piezo actuated mirrors for illumination pattern homogenization. Raw volumetric image deconvolution was done using the Richardson Lucy algorithm implementation in the flowdec Python package^71^ with 10 iterations.

